# Nutrition and exercise diabetes knowledge and practices of family members of patients in rural areas of Limpopo Province, South Africa

**DOI:** 10.1101/2022.07.27.501684

**Authors:** Mabitsela Mphasha, Linda Skaal, Tebogo Mothiba

## Abstract

Family Members (FMs) offers basic assistance with meals and exercise, both of which are critical in diabetes control. Support from family has been linked to better outcomes. Family support, on the other hand, might lead to poor diabetes outcomes, especially when FMs lack information. Patients’ outcomes can be influenced by established family practices. As a result, the goal of this study is to determine diabetes FMs’ knowledge and practices in the areas of nutrition and exercise. On 200 FMs selected via convenient sampling from rural clinics in Senwabarwana, Limpopo Province, a quantitative approach and cross-sectional descriptive design were used. Close-ended questionnaires were used to collect data, which were then analyzed using Statistical Package for Social Sciences Software v27.0 and descriptive and inferential statistical analysis. Only 31% of participants had great overall knowledge, according to the findings, and only 9% had good practice. Over half of the participants (53%) believe that overweight diabetes patients should skip meals in order to lose weight, and 3.5% and 19%, respectively, are familiar with exercise prescriptions and glucose measurements. Only 35,5% of people eat breakfast every day, whereas the overwhelming majority (87,5%) exercise. The findings of this study show that patient’s FMs need to enhance their diet and exercise diabetes care knowledge and practices. To reduce diabetes prevalence and its detrimental impact on patients’ diabetes treatment, a combined strategy of community-based awareness campaigns and a family-centered approach is proposed, as well as behavior change intervention.

## Introduction and background

The global burden of Diabetes Mellitus (DM) is rising, with 537 million persons aged 20 to 79 years living with the disease, with roughly 24 million adults living with the disease in Africa (International Diabetes Federation (IDF), 2021). Furthermore, 13 million persons in Africa have undiagnosed diabetes (IDF, 2021). South Africa’ s diabetes prevalence has risen over time, from 4.5 percent in 2010 to 12.7 percent in 2019. (Grundlingh et al., 2022). Diabetes rates are projected to rise due to sedentary lifestyles (Grundlingh et al., 2022), as well as a family history of the disease (Grundlingh et al., 2022). Genetic variables have been discovered to play a substantial role in diabetes prevalence in families (Ali, 2013). Due to a family history of Diabetes Mellitus, family members of diabetes patients are already at risk of developing the disease (Papazafiropoulou et al., 2017). Nutrition and exercise are crucial in diabetes management, and making changes in these areas may improve results and reduce diabetes prevalence (IDF, 2021). Obesity, which has been recognized as one of the key predisposing factors to diabetes, is caused by poor eating habits and a lack of physical activity (Ganu et al., 2016).

Knowledge, motivation, and individuals’ ability in acquiring, comprehending, appraising, and using health information are all linked to knowledge in health (He et al., 2016). Appropriate diabetes knowledge is linked to increased quality of life and the prevention of complications (Zowgar et al., 2018), and may aid in the management of diabetes, the prevention of complications, and the development of diabetes in those at risk (Alemayehu et al., 2020). On the other hand, a lack of information is associated to an increased risk of acquiring diabetes (Spronk et al., 2014). Poor diabetes knowledge also normalizes a lifestyle that greatly contributes to obesity, which is now recognized as a major risk factor for diabetes (Ganu et al., 2016). According to a Saudi Arabian study, methods for enhancing diabetes knowledge must be integrated into existing healthcare systems and processes in order to promote public awareness and empowerment (Alanazi et al., 2018). It has been stated that raising public awareness about diabetes, which has been found to be insufficient (Alemayehu et al., 2020), can help to minimize the chance of having diabetes. Knowledge alone does not guarantee the adoption of healthy behavior, according to Ajzen et al. (2011), just as ignorance is mostly responsible for ill behavior. Furthermore, combining information with motivation to improve behavior results in positive behavior change (Ajzen et al., 2011).

Health-related practices have been found to influence subjective perceptions of health, as well as self-rated health and overall quality of life (Maclean et al., 2004). Physical inactivity, as well as the consumption of fatty, sugary, and excessive portion sizes of food, were discovered to play a substantial role in the development of diabetes (Kyrou et al., 2020). In the face of social, physical, and emotional problems, health is defined as the ability to adapt and self-manage (Huber et al., 2011). Because family members of diabetes patients are already at risk of having diabetes due to a family history of the disease (Papazafiropoulou et al., 2017), it is critical that they adapt and self-manage in order to reduce the risk of developing diabetes. Similarly, established family practices may have an impact on a patient’ s diabetes care. In the management of T2DM and the prevention of diabetes, behavior change and intensive lifestyle interventions are critical (le Roux et al., 2019). Because family members provide home care for patients, family health practices are crucial.

The vast majority of everyday diabetes treatment takes place in patients’ homes, and Family Members (FMs) have been highlighted as a significant source of support for diabetic patients (Mphasha, Mothiba & Skaal, 2021). Despite research indicating that family members are expected to offer assistance and care at home, where diabetic patients receive the majority of their care, There is a lack of data including non-diabetic family members of people with diabetes, which creates a gap that this study aims to fill. Support from family is linked to better diabetes outcomes, quality of life, health status, and the prevention of diabetic complications (Mayberry & Osborn, 2012). Family support, on the other hand, can be harmful and contribute to poor diabetes outcomes, as well as a decline in health and quality of life, especially when FMs are insufficient or lack diabetes care understanding (Mphasha, Mothiba & Skaal, 2021). As a result, it’ s vital to measure diabetes care knowledge among FMs of diabetic patients. Patients’ diabetes results might be influenced negatively or favourably by family-established cultures, behaviors, and practices. As a result, it’ s critical to comprehend family members’ habits in order to comprehend how they may affect diabetes outcomes. This study aims to investigate diabetes care knowledge and practices among family members of patients in the areas of nutrition and exercise.

## Methodology

### Research Design

This study used a quantitative technique and a cross-sectional descriptive study design.

### Setting

The research was carried out in the rural Blouberg Municipality in the Senwabarwana area of the Capricorn District, Limpopo Province, South Africa.

### Population and sampling of the study

The participants in this study were non-diabetic family members of diabetes patients. A total of 200 people were chosen at the clinics utilizing convenient sampling. The sample size was determined using the overall statistics of diabetic patients, which were predicted to be 406. Yamane’ s (1967) formula for calculating sample size at 95 percent The formula for the confidence interval (CI) is: **n=N 1+N (e) 2; where n is the sample size, N is the population size (N=406), and e is the error margin (5%)**. Only family members of patients who have spent at least six months caring for them and are over the age of eighteen are eligible. Furthermore, family members with any chronic conditions were not included in the study.

### Instruments and data collection

A closed-ended questionnaire with three sections was used to collect data: demographic profile, knowledge and habits related to nutrition and exercise diabetes care. Eleven questions adapted from Muchiri, Rheeder, and Gericke (2017) and Le Roux et al. (2018) research using 3-point Likert scales “Yes, Not Sure; No” and “Regular; Sometimes; Never,” respectively, were used to develop knowledge and practices linked to dietary consumption. The questionnaire’ s reliability was guaranteed by piloting it in non-participating clinics, which yielded no changes. Peers were used to guarantee content validity (dietitians and supervisors).

### Data analysis

Quantitative data analysis was carried out by coding data and analyzing it with the Statistical Package for Social Sciences. Frequency distributions, means, and standard deviations were computed using descriptive statistics. The Chi-squared test with 95 percent confidence intervals was used to calculate relationships, with a p-value of 0.05 considered statistically significant.

### Consideration of ethical issues

The study was approved by the Turfloop Research Ethical Committee (TREC), which issued clearance certificate number TREC/35/2019: PG. The Limpopo Department of Health (DOH) approved the study with the reference number LP 201903-007. This research is part of a larger project, although this paper solely looked at diabetic dietary knowledge and its impact on intake. All of the participants signed a written informed consent form. The study was completely voluntary, and participants were advised of their freedom to withdraw at any time without penalty. The data of the participants was likewise kept private and confidential.

## Results

### Demographic profile of participants

Table 1 shows that the majority of participants (73%) were under the age of 60, were females (76%), had secondary or higher education (64%) and were just slightly more than half single (54%).

**Table 1:**
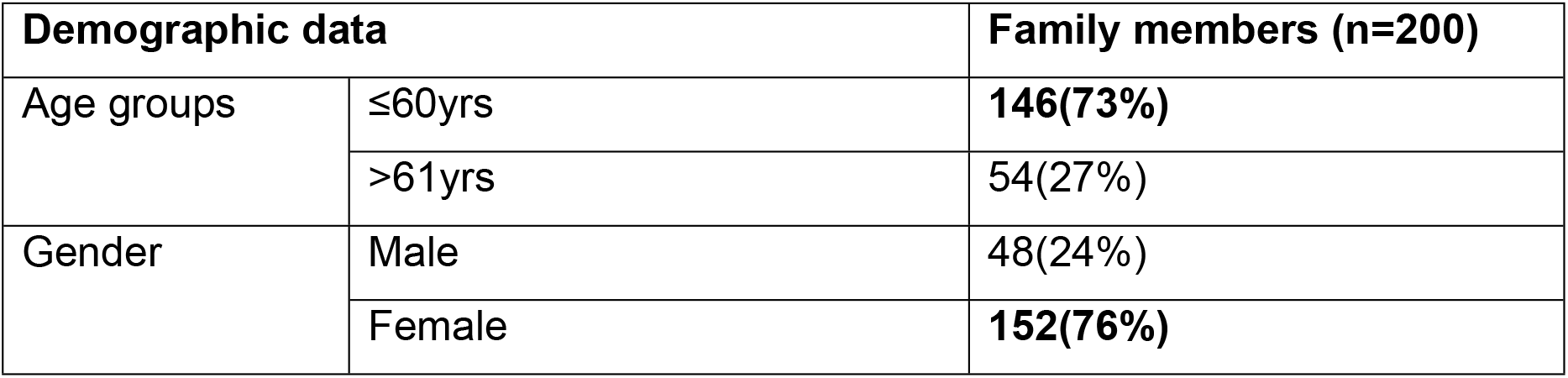
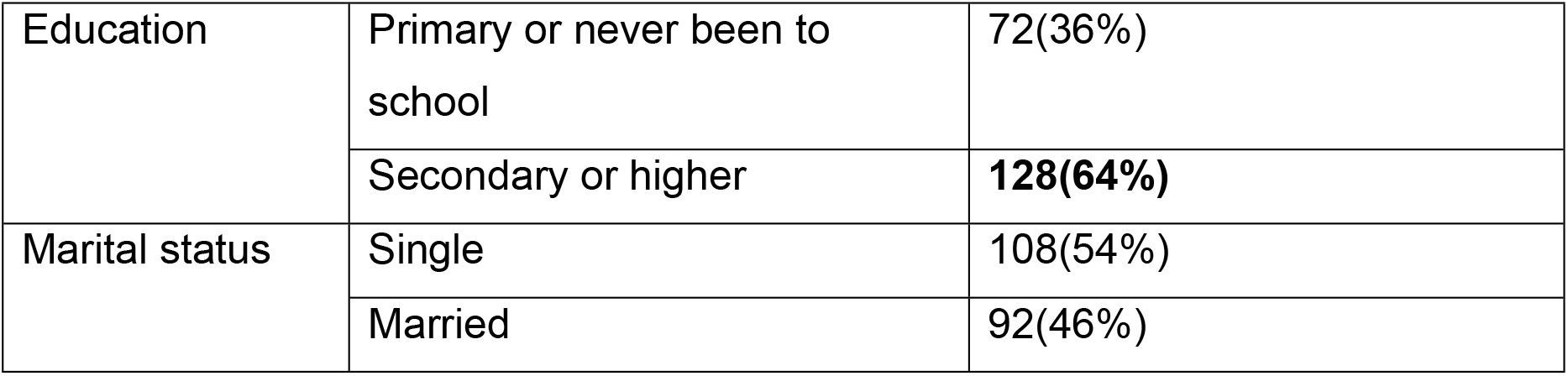
Demographic profile of participants.

| **Demographic data** |  | **Family members (n=200)** |
| --- | --- | --- |
| Age groups | ≤60yrs | **146(73%)** |
|  | >61yrs | 54(27%) |
| Gender | Male | 48(24%) |
|  | Female | **152(76%)** |
| Education | Primary or never been to school | 72(36%) |
|  | Secondary or higher | **128(64%)** |
| Marital status | Single | 108(54%) |
|  | Married | 92(46%) |

Figure 1 reveals that just about a third of the participants had outstanding knowledge (31%), followed by those who had good knowledge (24%), and those who had fair knowledge (22%).

**Fig 1:**
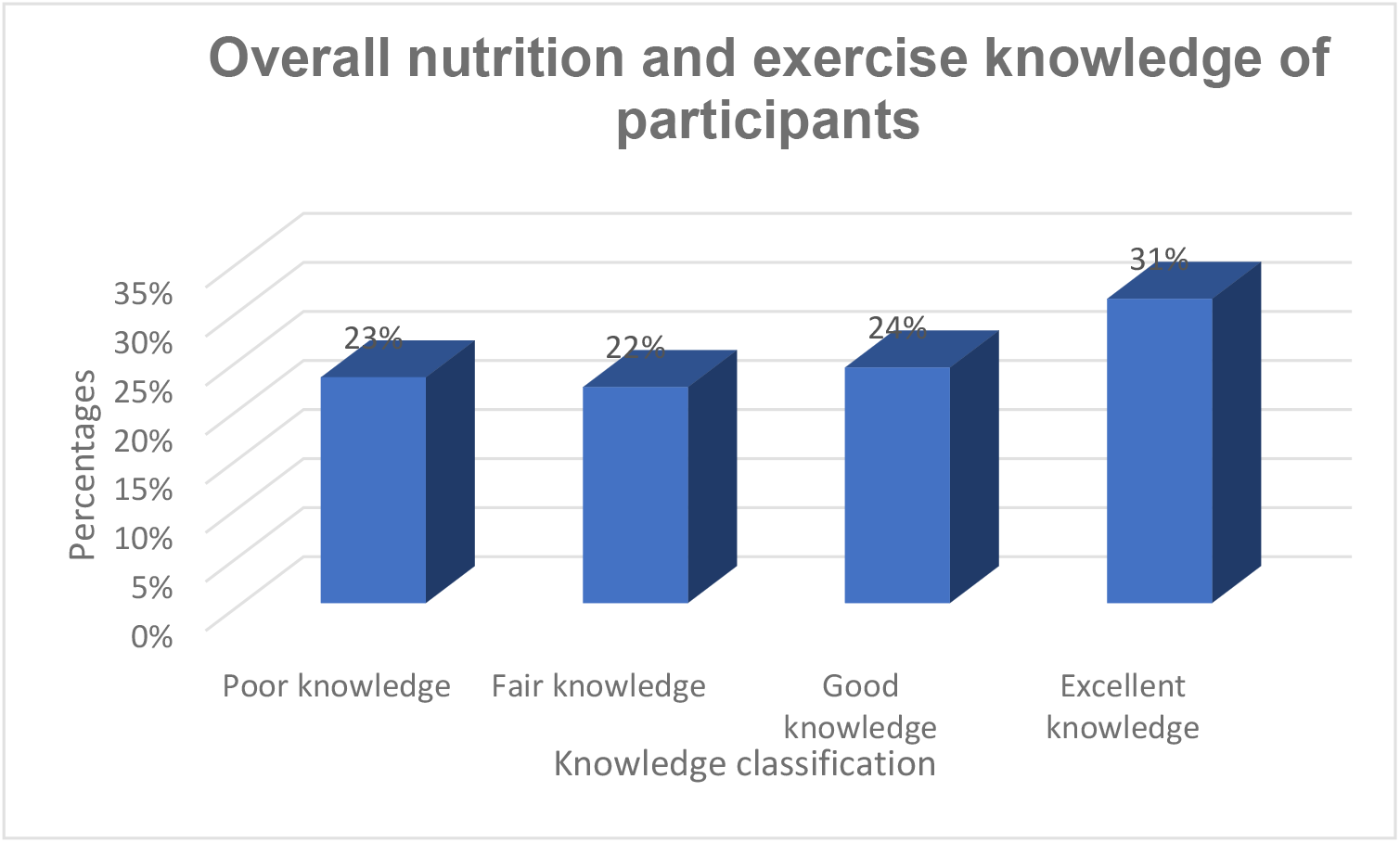
Overall nutrition and exercise diabetes care knowledge of participants.

Table 2 demonstrates that over half of the participants (53%) believe that overweight diabetics should miss meals in order to lose weight.

**Table 2:** Knowledge of participants regarding diabetes care through nutrition, % in rows; n=200.

| Knowledge of participants regarding diabetes care through nutrition | Yes | Not sure | No |
| --- | --- | --- | --- |
| Nutrition plays an important part in diabetes management. | 181(90,5%) | 1(0,5%) | 18(9%) |
| Fruits and vegetables must be eaten because they are good in managing blood sugar. | 177(88,5%) | 4(2%) | 19(9,5%) |
| Diabetes patients should eat small, frequent meals regularly to manage blood sugar. | 162(81%) | 4(2%) | 34(17%) |
| Whole-grains high in fiber are recommended as a healthy source of carbohydrate for diabetes patients. | 153(76,5%) | 5(2,5%) | 42(21%) |
| *It is recommended that diabetes patient who is overweight should skip meals to lose weight.* | 106(53%) | 10(5%) | 84(42%) |
| When preparing meat for diabetes patients, it is recommended to remove visible fats from red meat, and also to eat chicken without skin. | 128(64%) | 2(1%) | 70(35%) |
| If a diabetes patient eat large portion size of food at once may lead to increased blood sugar. | 129(64,5%) | 3(1,5%) | 68(34%) |
| Diabetes patients should avoid high fat dairy products including high animal proteins must be avoided. | 118(59%) | 5(2,5%) | 77(38,5%) |
| It is good for diabetes patients to cut back on salty food including high sodium food such as processed food. | 140(70%) | 4(2%) | 56(28%) |
| It is good for diabetes patients to cut back on sugary food including avoiding added sugar in drinks and food. | 143(71,5%) | 6(3%) | 51(25,5%) |
| Diabetes patients should avoid fried food and other food high in fats. | 147(73,5%) | 3(1,5%) | 50(25%) |

Table 3 demonstrates that the majority of participants (89.5%) are unaware of exercise prescriptions or tolerable blood glucose limits (75,5%).

**Table 3:** Knowledge of participants regarding diabetes care through exercise, % in rows; n=200.

| Knowledge of caregivers regarding diabetes care through exercise | Yes | Not sure | No |
| --- | --- | --- | --- |
| Exercise is important in diabetes management and help control blood sugar | 170(85%) | 3(1,5%) | 27(13,5%) |
| When a diabetes patient exercises blood sugar, blood pressure and cholesterol levels stay on track. | 167(83,5%) | 2(1%) | 31(15,5%) |
| Exercising can help diabetes patients to prevent and/or reduce diabetes-related health problems or complications. | 162(81%) | 6(3%) | 32(16%) |
| It is recommended that a diabetes patient should stop exercising, in the event of feeling dizziness, shortness of breath and pains | 148(74%) | 5(2,5%) | 47(23,5%) |
| When a diabetes patient engages in prolonged exercise, it is encouraged to eat before and after exercise, including drinking fast acting carbohydrate drinks during exercise. | 151(75,5%) | 3(1,5%) | 46(23%) |
| It is recommended for diabetes patient to check blood sugar before and after exercise. | 150(75%) | 4(2%) | 46(23%) |
| A diabetes patient should wait until blood sugar becomes normal when he/she discovers that his/her blood sugar is high just before exercise. | 135(67,5%) | 10(5%) | 55(27,5%) |
| *It is recommended that a diabetes patient should at least exercise for a minimum amount of 30 minutes, 3 days in a week.* | 7(3,5%) | 14(7%) | 179(89,5%) |
| *When a diabetes patient who is on insulin and does running exercises, should not inject themselves on thighs.* | 6(3%) | 10(5%) | 184(92%) |
| *The amount of insulin in the body decreases during exercise.* | 48(24%) | 13(6,5%) | 139(69,5%) |
| *The acceptable blood glucose levels of a diabetes patients is a range of 4-8mmol/L.* | 38(19%) | 11(5,5%) | 151(75,5%) |

Figure 2 demonstrates that nearly half of the participants (48%) had fair practice, followed by 43 %who had bad practice, and only 9% who had good practice.

**Fig 2:**
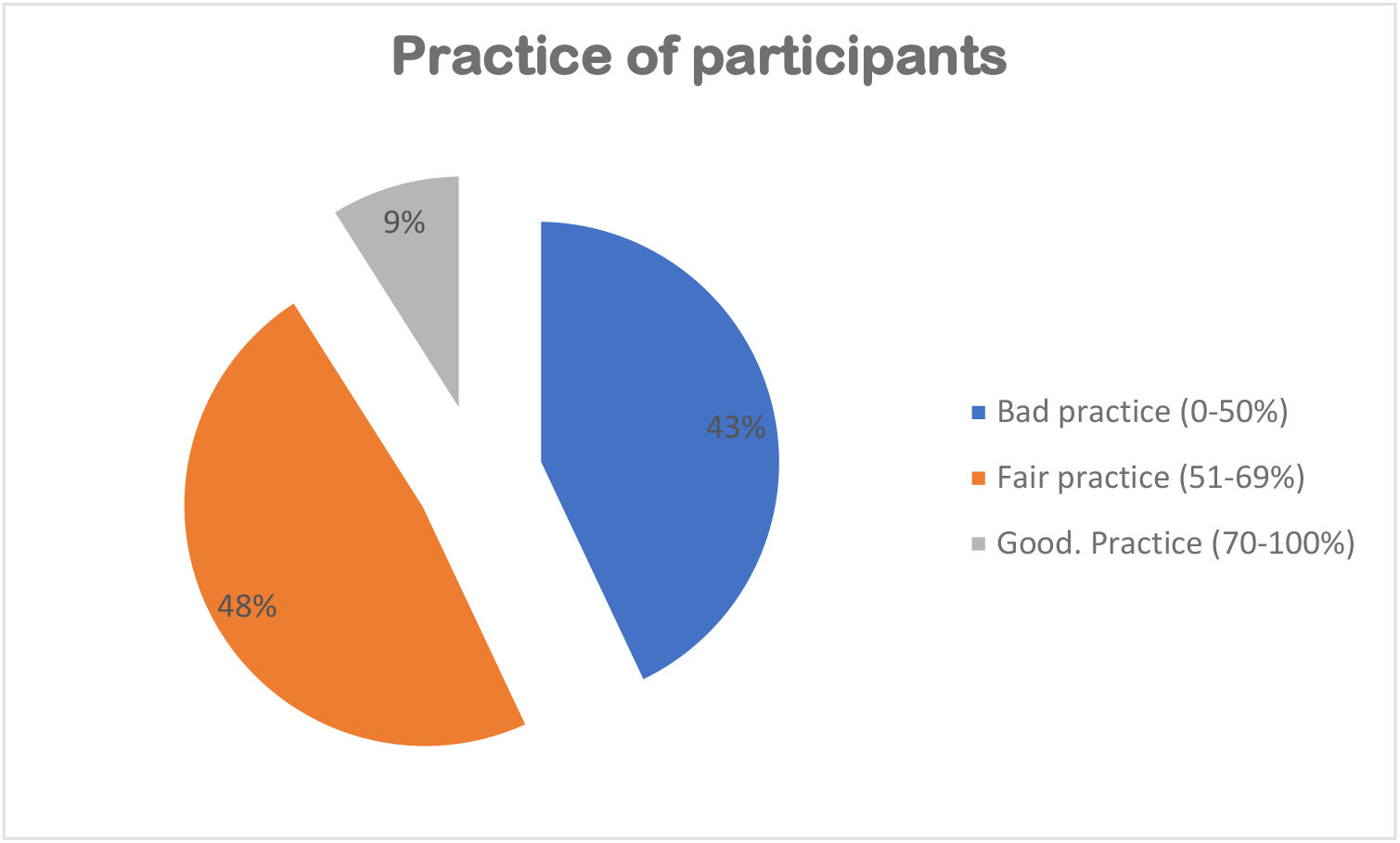
Overall practice of participants.

Table 4 shows that only 35,5% of over a third of participants eat breakfast (35,5%) and also eat meat, chicken, fish, mopani-worms, eggs, and milk (38,5%) regularly.

**Table 4:** Practice of participants regarding diabetes management through nutrition, % in rows; n=200.

| Practice of participants regarding diabetes care through nutrition | Regularly (4 or more times a week) | Sometimes (1-3 times a week) | Never |
| --- | --- | --- | --- |
| How often did you have breakfast in the last week? | 71(35,5%) | 119(59,5%) | 10(5%) |
| How often did you have lunch and dinner in the last week? | 138(69%) | 62(31%) | 0(0%) |
| How frequently do you consume fried and fatty foods? | 16(8%) | 127(63,5%) | 57(28,5%) |
| How often do you consume sugary foods and beverages? | 10(5%) | 129(64,5%) | 61(30,5%) |
| How often do you consume alcoholic beverages? | 2(1%) | 50(25%) | 148(74%) |
| How frequently do you consume vegetables? | 36(18%) | 155(77,5%) | 9(4,5%) |
| How frequently do you consume fruits? | 34(17%) | 160(80%) | 6(3%) |
| How often do you eat meat, chicken, fish, mopani-worms, eggs and milk | 77(38,5%) | 115(57,5%) | 8(4%) |
| How often eat high fiber food such as beans, lentils, legumes and wholegrain foods among others? | 58(29%) | 136(68%) | 6(3%) |
| How frequently do you eat starchy food such as porridge, rice, samp and bread? | 188(94%) | 12(6%) | 0(0%) |
| How often do you eat salty food such as processed food like polony or added salt? | 17(8,5%) | 120(60%) | 63(31,5%) |

Table 5 shows that overwhelming majority of participants exercises (87%). Only 32% exercised for at least 30 minutes on three days per week in the last month.

**Table 5:**
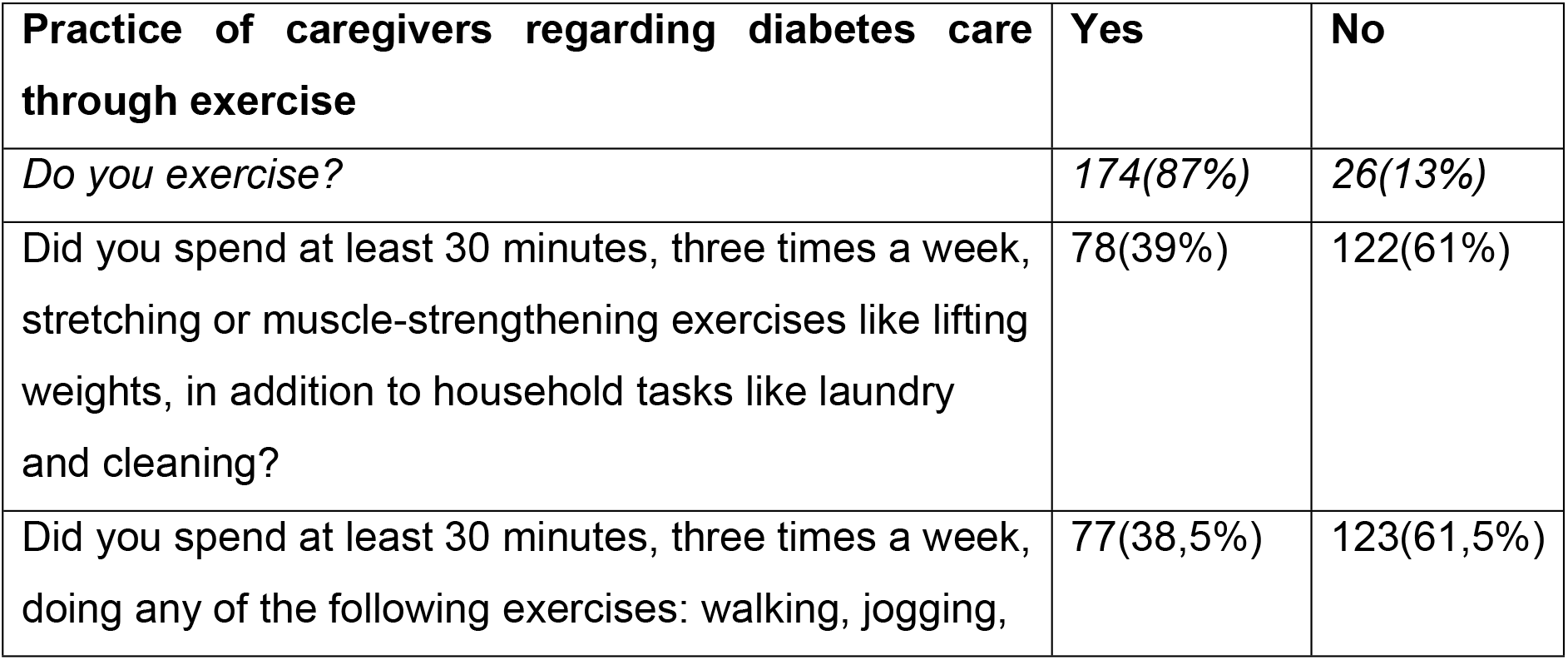
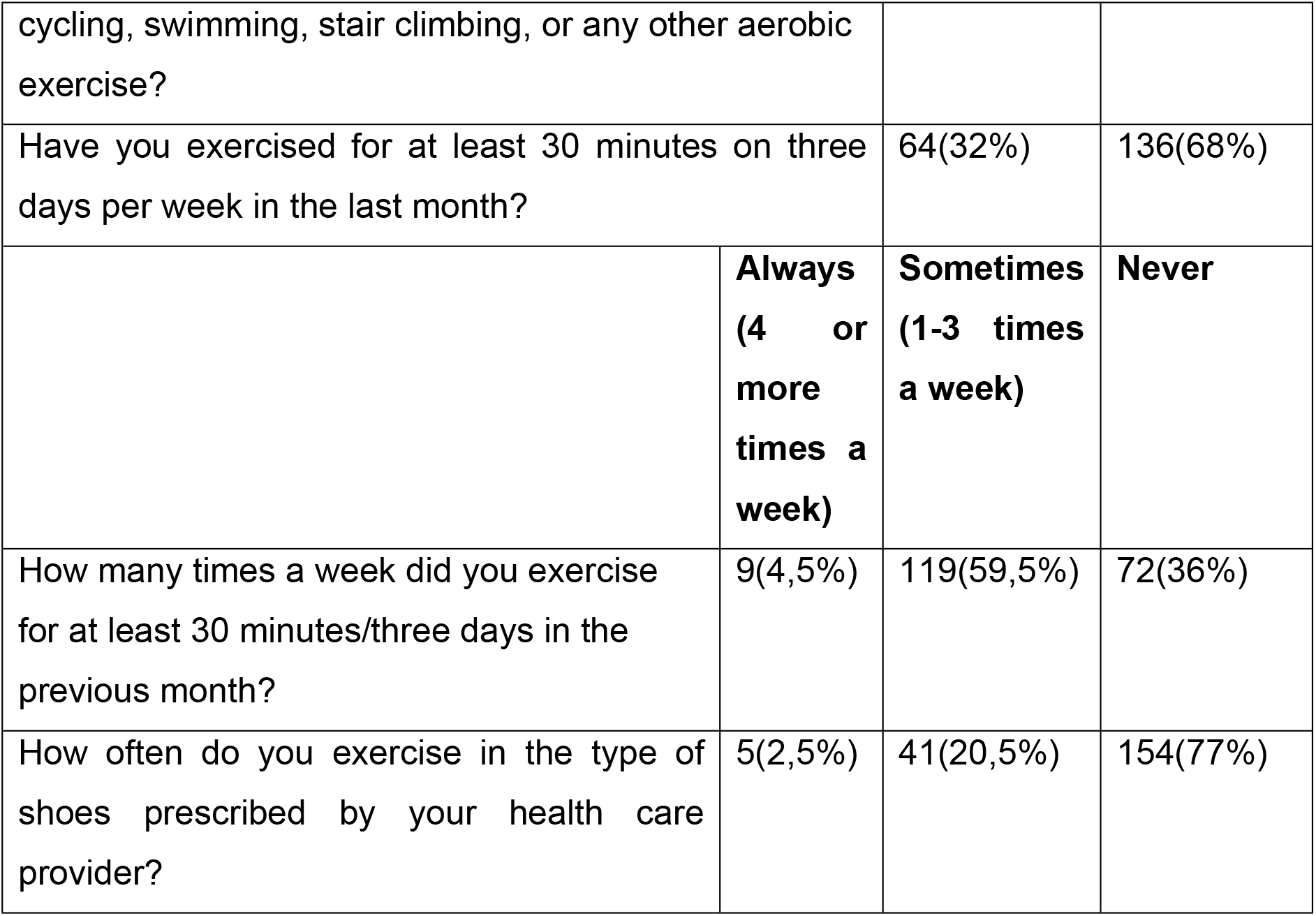
Practice of participants regarding diabetes management through exercise, % in rows; n=200.

## Discussions

This study recognizes that family members of diabetic patients are an important source of care and support for patients. Support from family is associated to better diabetes results, but it can also lead to complications and bad outcomes. As a result, the goal of this study was to find out how well diabetes patients’ family members knew about nutrition and exercise.

According to the findings of this study, 31% of participants had great understanding of total nutrition and exercise diabetes management. This was in line with the findings of an Ethiopian community-based cross-sectional study, which found that non-diabetic community members had little knowledge of diabetes (Alemayehu, Dagne, & Dagnew, 2020). Family members with good diabetes knowledge and its serious complications may be able to assist family members with diabetes in taking appropriate measures to control the disease, such as keeping medical visits and adhering to lifestyle adjustments (Shrivastava et al., 2013). As a result, participants in this study may not be able to provide adequate support to patients in order to help them achieve better diabetes results. There is an immediate need to increase nutrition and exercise diabetes care knowledge in order to enable them to understand the dos and don’ ts of diabetes care in order to protect patients’ health. Given that they are already at risk, empowering family members will also assist them in initiating lifestyle changes to decrease or prevent diabetes (Adu et al., 2019). Individuals who have sufficient knowledge may be motivated or persuaded to live a healthy lifestyle (Ajzen et al., 2011).

Over a third of the participants in this survey were unaware that eating large portions is bad for diabetes patients, nearly half were unaware that high-fat dairy products should be avoided, and over half believed that overweight diabetes patients should skip meals to lose weight. This research backs up a Kenyan community-based cross-sectional study that found inadequate diabetes dietary awareness among non-diabetic family members of patients (Kiberenge et al., 2010). Diabetic patients’ family members are supposed to help them. Obesity may develop as a result of the consumption of high portion sizes and fatty meals (Rolls, 2014). Obesity has a key role in diabetes incidence and complications among patients (Ganu et al., 2016), as well as raising the risk of diabetes among patients’ family members due to a family history of diabetes (Papazafiropoulou et al., 2017). Only 35.5 percent of individuals eat breakfast every day, and more than two-thirds eat fatty (63.5%), sugary (64,5%), and salty (60%) items one to three times each week. Over two-thirds of participants (69%) said they eat both lunch and dinner on a regular basis, contradicting reports that the majority of people do not eat both lunch and dinner due to cost constraints (Dwyer, 2014). Increased fat and oil intake has been reported in South Africa and other developing countries, according to this study (Popkin, 2004). This established family practice may have a negative impact on diabetes treatment for diabetic family members, as well as increasing the likelihood of family members obtaining diabetes.

Participants in this study understand the importance of exercise in diabetes control. However, this does not guarantee that participants will urge diabetes patients to exercise, as information is not enough to lead to an active lifestyle (Ajzen et al., 2011). Physical activity in rural areas has been found to be influenced by factors such as urbanization (Aristides et al., 2014). Physical activity is linked to improved diabetes outcomes in patients (Cannata et al., 2020) and lower diabetes risk in vulnerable people (Hjerkind et al., 2017). The majority of participants are aware that checking glucose levels before and after exercise is beneficial for diabetics, but they lack knowledge about exercise prescriptions and glucose readings. The vast majority of participants in this survey (87 percent) stated that they exercise, which can be attributed to their understanding of the importance of exercise. However, only 4,5 percent of individuals said they had exercised in the previous week, raising questions about consistency. As a result, it is necessary to reinforce exercise through a behavior modification method in order to make it a habit for people to exercise regularly and sustainably.

Through a family-centered care approach, it is critical to increase family members’ nutrition and exercise diabetes care knowledge and habits. This method will allow family members of diabetic patients to collaborate and participate in their care. As a result, family members will be empowered and equipped with the dos and don’ ts of diabetes care in order to improve outcomes and reduce diabetes prevalence. Diabetes programs centered in the community are also recommended for raising public awareness and encouraging people to live a healthy lifestyle in order to prevent diseases like diabetes.

## Conclusion

Family members of diabetic patients have little knowledge and poor practices when it comes to diabetes nutrition and activity. As a result, family members’ support of patients may be counterproductive and have a negative impact on results. Furthermore, due to family history, insufficient knowledge and poor practices related to nutrition and exercise, which may contribute to obesity, family members are at an increased risk of acquiring diabetes. To reduce diabetes prevalence and its detrimental impact on patients’ diabetes treatment, a combined strategy of community-based awareness campaigns and a family-centered approach is proposed, as well as behavior change intervention.

### Recommendations

In diabetes management, a combined intervention of community awareness and family-centered care should be implemented.

Behavioural change is supported to enhance nutrition and exercise practices, resulting in better diabetes management and outcomes for patients.

### Limitations

Participants in this study were family members who accompanied their diabetic family member to the clinic over the study’ s duration.

## Authors contribution

Mphasha was a project leader in charge of data gathering and interpretation, and he contributed 50% of the article’ s authoring. Skaal analyzed data and oversaw data collecting and interpretation; she was responsible for 40% of the article’ s authoring. Mothiba assisted with data collection and interpretation and contributed 10% to the article’ s composition. All of the authors signed off on the final manuscript.

## Availability of data and materials

The information in this article comes from diabetes patients and their non-diabetic family members in the Senwabarwana district of Limpopo Province, South Africa. Because more publications are anticipated, the dataset generated or analyzed during this study is not publicly available, although it can be obtained from the corresponding author.

## Disclaimer

The opinions expressed in the articles are those of the authors and do not reflect the institution’ s official position.

